# Screening and potent applicability analysis of commonly used pesticides against desert locust: an integrative entomo-informatics approach

**DOI:** 10.1101/2020.10.19.345595

**Authors:** Anik Banik, Md. Fuad Mondal, Md. Mostafigur Rahman Khan, Sheikh Rashel Ahmed, Md. Mehedi Hasan

**Affiliations:** Faculty of Biotechnology and Genetic Engineering, Sylhet Agricultural University, Sylhet-3100, Bangladesh; Department of Entomology, Sylhet Agricultural University, Sylhet-3100, Bangladesh; Department of Plant and Environmental Biotechnology, Sylhet Agricultural University, Sylhet-3100, Bangladesh

**Keywords:** Desert locusts, GIS mapping, pesticides, phylogenic analysis, molecular docking, microbial volatile compound, molecular dynamics

## Abstract

The locust problem is a global threat for food security. Locusts can fly and migrate overseas within a zip and creating a large-scale devastation to the diversified agro-ecosystem. GIS based analysis showed the recent movement of locusts, among them *Schistocerca gregaria* and *Locusta migratoria* are predominant in Indian subcontinent and are found more notorious and devastating one. This devastation needs to be stopped to save human race from food deprivation. In our study, we screened some commonly used agricultural pesticides and strongly recommended three of them *viz*. biphenthrin, diafenthiuron and silafluofen which might be potential to control the desert locusts based on their binding affinity towards the locust’s survival proteins. Our phylogenetic analysis reveals that these three recommended pesticides might also show potency to the other locust species as well as they are also way safer than the other commercially available pesticides. These proposed pesticide’s bioactive analogs from fungus and bacteria may also show efficacy as next generation controlling measures of locust as well as different kind of pests. These recommended pesticides are expected to be highly effective against locusts and needs to bring forward by the entomologists’ by performing experimental field trials.

**Highlights:**

1. GIS map unmasked the 2020 migratory pattern of locusts which now predominant towards Indian subcontinent.
2. Biphenthrin, diafenthiuron and silafluofen showed maximum binding affinity.
3. Biphenthrin and diafenthiuron were relatively safer than silafluofen.
4. Bioactive analogs from fungus and bacteria could be an alternative to control locusts.
5. Pesticides inhibition hotspots for desert locusts were unrevealed.

## 1. Introduction

Locusts belongs to the family Acrididae, are the short-horned grasshoppers which are one of the most diverse agricultural and ecologically important pests [1,2]. There are 19 locust species reported around the world by different study [3–5]. Among them, some species *viz. Locusta migratoria, Schistocerca gragaria, Locustana pardalina, Schistocerca piceifrons, Calliptamus italicus, Dociostaurus maroccanus, Nomadacris septemfasciata, Schistocerca cancellata* are devastating. Moreover, *L. migratoria* and *S. gragaria* are most devastating in recent year [6,7]. The solitary locust become more abundant by changing their behavior under certain circumstances, becoming gregarious and can be serious pests of agriculture due to their ability to form dense and highly mobile swarms [2,8]. Gregarious locusts in one region can migrate to another region in a single night about hundreds of kilometers and plagues can spread out continents [3].

These divesting species affected more than 500 plant species including different crops, fruits and trees [2,9]. An adult desert locust can consume its own weight fresh food per day and 1 km^2^ size swarm desert locusts eat about 35,000 people’s food per day [9]. Moreover, the species of locusts was a reason for a devastating plague in 2470 to 2220 BC was mentioned in the Holy Book of Bible taking place in Egypt [10,11]. In addition, the migratory locust from China named Asian migratory locust showed their devastating phenomenon as plague from 200 BC, which was accelerated by droughts and flood events [12,13] Different Mediterranean and sub-Saharan country as well as drought area of Africa, Europe, middle Asia and eastern Asia were affected by locust several times [14–16]. The incidence of 1949-57, Morocco lost $60 million and in 1958 Ethiopia lost 167,000 MT of grain. At the end of last century, $310 million was used globally for only locust control [6].

The current outbreak began in 2018 when Cyclone Mekunu produced heavy rains in the Rub’ al Khali of the Arabian Peninsula. In spring 2019, swarms spread from these areas, and by June 2019, the locusts spread north to Iran, Pakistan, India and south to East Africa, particularly the Horn of Africa [17]. According to the Food and Agriculture Organization (FAO), the desert locust is spreading over more than 30 countries in the world and reported as most dangerous migratory pest. In this year, locusts swarm already affected large numbers in several countries, including Ethiopia, Eritrea, Yemen, Oman, Uganda, Kenya, Somalia, Iran, Saudi Arabia, India and Pakistan. Recently, FAO forecasted that, explosive multiplication of Locust swarms’ forms disasters for large parts of Asia and Africa in this year.

Fight against locust using pesticides is a common practice in many countries of the world. In addition, fight against these notorious locust different biological control approaches are also being used [7,18]. The affected countries are using different group of pesticides in large scale but not yet much effective as their expectation [6]. As locust invade with a huge swarm it’s very difficult to control [6]. Moreover, the migratory behavior also affects the control process. Therefore, countries are facing challenges for controlling thus pest. In such a circumstances computational approach could be better way to find effective solution for managing this notorious pest. Computational chemistry in particular, virtual screening can provide valuable insights in hit and lead effective compound discovery. Numerous software tools have been developed to meet the purpose [19]. In this study, we evaluated several commercially available pesticides to find the best fitted pesticide among them to combat the devastating notorious locust by using several computational approaches.

## 2. Materials and Methods

### 2.1 GIS based mapping of Desert locust infestation areas

The GIS based data of Desert locust infestation point and infestation area were retrieved from the “Locust Hub” and “Locust Watch” sorting according to the period of infestation and analyzed by the different analyzing tools of GIS base software (ArcGIS 10.7.1) (https://locust-hub-hqfao.hub.arcgis.com/, http://www.fao.org/ag/locusts/en/info/info/index.html). The retrieved data period was June, 1985 to September, 2020.

### 2.1. Retrieval of target locust proteins/protein-domains and insecticides

The RCSB Protein Data Bank [20] was used for the retrieval of 3D structures of *S. gregaria* lipid binding protein (PDB ID: 2GVS) and Muscle fatty acid binding protein (PDB ID: 1FTP) along with *L. migratoria* fatty acid binding protein from flight muscle (PDB ID: 2FLJ) and Odorant binding protein (PDB ID: 4PT1). A total of 31 insecticide (Supplementary file 1), used against different orthoptera (e.g. grasshopper locust, cricket etc.) were extracted from PubChem database (https://pubchem.ncbi.nlm.nih.gov/) [21] in SDS (3D) format (Table 1). Now the OpenBabel v3.1.1 software was utilized to convert the retrieved SDF structure into PDB format for further analysis [22,23].

**Table 1:** Group name, Agrochemical category, Mode of action and Impact Area of available commercial insecticides (IRAC, 2020)

| **Insecticides Name** | **PubChem ID** | **Group Name** | **Agrochemical Category** | **Mode of Mechanism** | **Impact Area** |
| --- | --- | --- | --- | --- | --- |
| Aldicarb | 9570071 | Carbamates | Nematicides, Insecticides, Acaricides | Acetylcholinesterase (AChE) inhibitors | Nerve |
| Aldrin | 12310947 | Organochlorines | Insecticides | GABA-gated chloride channel blockers | Nerve |
| Allethrin | 11442 | Pyrethroids | Insecticides | Sodium channel modulators | Nerve |
| Azamethiphos | 71482 | Organophosphates | Insecticides | Acetylcholinesterase (AChE) inhibitors | Nerve |
| Biphenethrin | 6442842 | Pyrethroids | Insecticides, Acaricides | Sodium channel modulators | Nerve |
| Carbaryl | 6129 | Carbamates | Insecticides, Plant growth regulators | Acetylcholinesterase (AChE) inhibitors | Nerve |
| Carbendazim | 25429 | Carbamates | Alkaline solution, Insecticide | Acetylcholinesterase (AChE) inhibitors | Nerve |
| Carbofuran | 2566 | Carbamates | Nematicides, Transformation products, Acaricides, Insecticides | Acetylcholinesterase (AChE) inhibitors | Nerve |
| Chloropyriphos | 2730 | Organophosphates | Insecticides, Acaricides | Acetylcholinesterase (AChE) inhibitors | Nerve |
| Cyfluthrin | 104926 | Pyrethroids | Insecticides, Acaricides | Sodium channel modulators | Nerve |
| Cypermethrin | 2912 | Pyrethroids | Insecticides, Acaricides | Sodium channel modulators | Nerve |
| Diafenthiuron | 3034380 | Organosulfur | Miticides, Acaricides, Insecticides | Inhibitors of mitochondrial ATP synthase | Energy metabolism |
| Diazinon | 3017 | Organophosphates | Repellents, Veterinary substances, Acaricides, Insecticides | Acetylcholinesterase (AChE) inhibitors | Nerve |
| Dichlorvos | 3039 | Organophosphates | Transformation products, Acaricides, Insecticides | Acetylcholinesterase (AChE) inhibitors | Nerve |
| Dieldrin | 969491 | Organochlorines | Insecticide | GABA-gated chloride channel blockers | Nerve |
| Dimethoate | 3082 | Organophosphates | Transformation products, Acaricides, Insecticides | Acetylcholinesterase (AChE) inhibitors | Nerve |
| Dinotefuran | 197701 | Neonicotinoids | Insecticides | Nicotinic acetylcholine receptor (nAChR) competitive modulators | Nerve |
| Endrin | 12358480 | Organochlorines | Insecticide, rodenticides | GABA-gated chloride channel blockers | Nerve |
| Ethienocarb | 73555894 | Carbamates | Insecticides | Acetylcholinesterase (AChE) inhibitors | Nerve |
| Etofenprox | 71245 | Pyrethroids | Insecticides | Sodium channel modulators | Nerve |
| Fenitrothion | 31200 | Organophosphates | Insecticides, Acaricides | Acetylcholinesterase (AChE) inhibitors | Nerve |
| Fenobucarb | 19588 | Carbamates | Insecticides | Acetylcholinesterase (AChE) inhibitors | Nerve |
| Fipronil | 3352 | Phenylpyrazoles | Insecticides | GABA-gated chloride channel blockers | Nerve |
| Heptachlor | 3589 | Organochlorines | Insecticides | GABA-gated chloride channel blockers | Nerve |
| Malathion | 4004 | Organophosphates | Insecticides, Acaricides | Acetylcholinesterase (AChE) inhibitors | Nerve |
| Methomyl | 5353758 | Carbamates | Insecticides | Acetylcholinesterase (AChE) inhibitors | Nerve |
| Methyl parathion | 4130 | Organophosphates | Insecticides, Repellants | Acetylcholinesterase (AChE) inhibitors | Nerve |
| Oxamyl | 9595287 | Carbamates | Insecticides, Nematicides | Acetylcholinesterase (AChE) inhibitors | Nerve |
| Parathion | 991 | Organophosphates | Insecticides, Acaricides | Acetylcholinesterase (AChE) inhibitors | Nerve |
| Phosmet | 12901 | Organophosphates | Insecticides | Acetylcholinesterase (AChE) inhibitors | Nerve |
| Silafluofen | 92430 | Pyrethroids | Insecticides | Sodium channel modulators | Nerve |

### 2.2. Phylogenic Analysis of proteins

Phylogenetic analysis is the means by which evolutionary relationships are calculated. The sequence of a particular gene or protein can be used in molecular phylogenetic research to determine the evolutionary relationships of organisms. Our targeted two species’s protein’s FASTA format were retrieved from PDB [20] and for homologous protein we performed BLASTp (https://blast.ncbi.nlm.nih.gov/Blast.cgi?PAGE=Proteins) excluding the parent organism. Then the tree was constructed by the maximum-likelihood method using MEGA X software [24].

### 2.3. Molecular docking of insecticides against locust proteins/protein-domains

Molecular docking is a key tool in structural molecular biology and it proposes structural hypotheses of how the ligands inhibit the target, which is invaluable in lead optimization [25]. The binding affinity of protein-ligand complexes were ranked via this effective method [26,27]. The binding affinity of 31 insecticides with different locust proteins/protein domains was calculated using PatchDock server which implies the shape complementary principal-based algorithm and the docking was done with clustering RMSD 4.0 and other default parameters [23,28,29]. For refining the protein-ligand docked complexes, FireDock refinement tool [30] was utilized. Finally, the visualization of protein-ligand complexes was visualized by Discovery Studio v3.1 [31] and PyMOL v2.0 [32].

### 2.4. Free binding energy calculation and Molecular Dynamics Stimulation

The LARMD server (http:/chemyang.ccnu.edu.cn/ccb/server/LARMD) was used to investigate the ligand-protein interactional binding mode based on conventional molecular dynamics [33]. The binding free energy (Δ*G* bind) was calculated based on binding energy (Δ*E* bind), solvation entropy (*T*Δ*S* sol) and conformational entropy (–*T*Δ*S*conf) [34]. The enthalpy and the entropy were calculated by the MM/GBSA method [35] and empirical method, respectively [36,37].

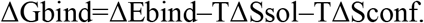

Molecular dynamics study was performed to signify protein-ligand complex stability for strengthening our prophesy. The normal modes of proteins can be compared to essential dynamics to determine their stability with their fluctuation plot analysis [38–40]. iMODS an online server, utilizes to analyze of normal modes (NMA) in internal coordinates which comprises deformability, B factor and Eigen value parameters which helps to understand the collective motion of proteins [41]. Further, stimulation was performed for 4ns on LARMD server to calculate the root mean square deviation (RMSD) and radius of gyration (Rg) of ligands and Root mean square fluctuation (RMSF). All the snapshots from the trajectory with a time interval per picosecond by using the Cpptraj module in the AMBER16 program were provided [42]. RMSD, Rg, and RMSF were simultaneously analyzed through this module.

### 2.5. Toxicity analysis of pesticides

The relative toxicity of top pesticide candidates was predicted via pkCSM web server. The server pkCSM, a novel method for predicting and optimizing small-molecule toxicity properties which relies on distance-based graph signatures [43]. Notably, the data mining process utilized for bird toxicity, rat toxicity, Fish toxicity of the pesticides through literature study [21,44,45].

### 2.6. Prediction of structural Analogs of pesticides

mVOC v2.0 web tools were used to identify potential small molecules against locust based on homology screening of predicted top pesticides. The server allowed ligand-based virtual screening of several libraries of small molecules to find available similar microbial volatile compound with known fungi and bacterial which are volatile emitters using several methods [46].

## 3. Results

### 3.1 GIS based mapping of Desert locust infestation areas

All the information of locust infestation showed the affected areas of Middle East, Northern Africa, Central and Southern Asia (Figure :1A). However, the recent outbreak of locust showed little bit different movement status. The new invasion area is gradually increasing in Southern Asia than the African region. The desert locust (*S. gregaria* and *L. migratoria*; most devastating in Middle East) moved to Indian subcontinent from Middle East over the Arab sea with long flight (Figure 1B). The results showed that the Ethiopia, Eritrea, Somalia, Yemen, Oman, Saudi Arabia, Pakistan are highly infested country among the desert locust infestation recorded country in 2020 outbreak (Figure 1:C). Moreover, several states of India also affected and faced threat in different crops (grain, pulse, vegetables and economically important trees) producing state. Furthermore, the neighbor country of Mali, Nigeria, Mauritania are challenging threat for desert locust rapid invasion.

**Figure 1.**
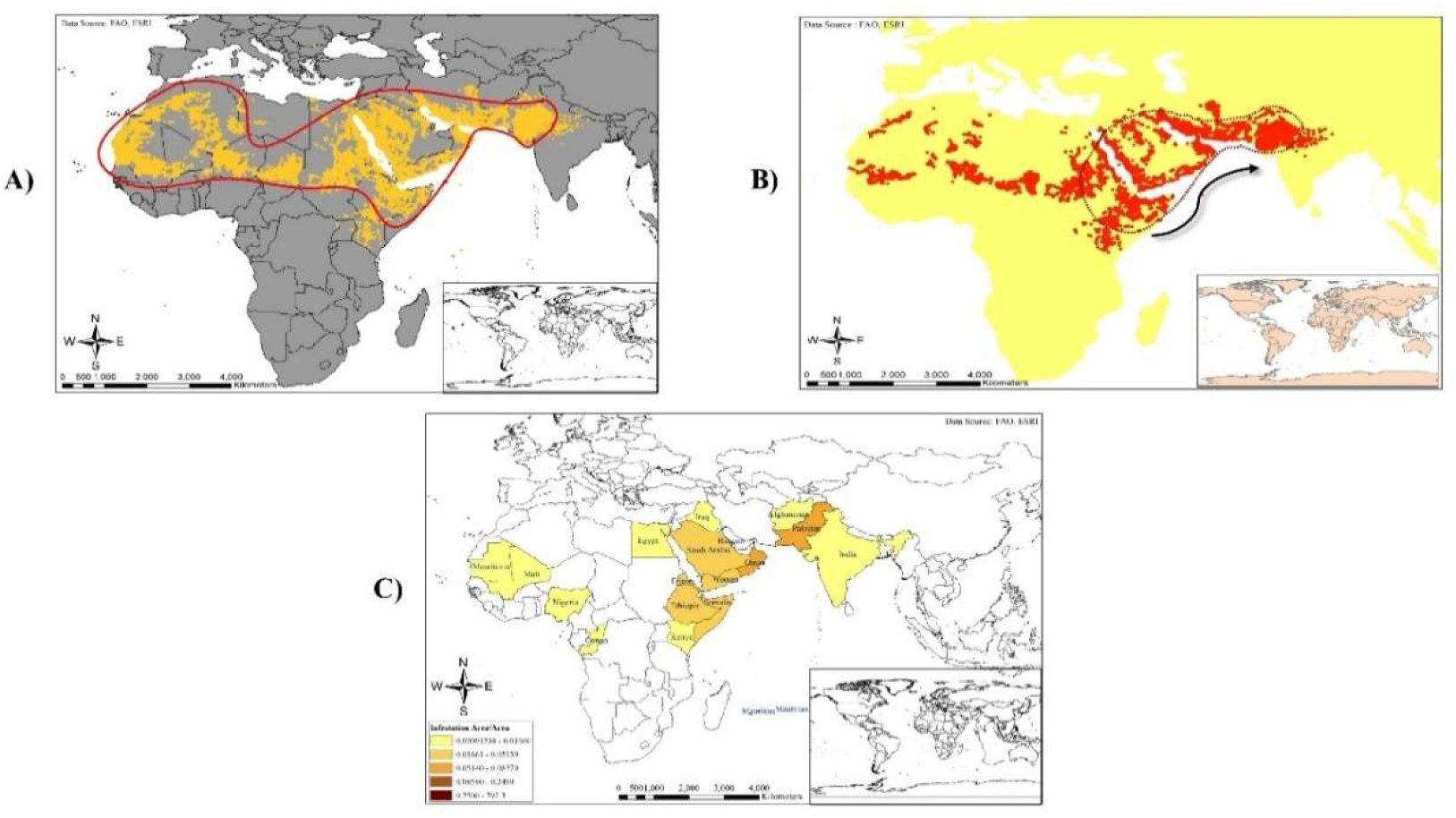
GIS based locust hotspots; A) Desert Locust infestation area over the world from June, 1985 to September, 2020; B) Desert Locust infestation area over the world in 2020; C) Desert Locust infestation severity in 2020 over the world.

### 3.2. Phylogenic Analysis of proteins

The phylogenetic analysis revealed that our targeted proteins from *S. gregaria* and *L. migratoria* are very much closely related to other locust species. Proteins like the *muscle fatty acid binding protein of S. gregaria* was closely related to several locust species protein such as *Halyomorpha haly, Riptortus pedestris, Nilaparvata lugens etc.* The CSPsg4 lipid binding protein from *S. gregaria* were closely related to proteins of *Ceracris kiangsu, Oedaleus infernalis, L. migratoria etc.* The fatty acid binding protein and the odorant binding protein from *L. migratoria* showed the close relationships with the *S. gregaria, H. haly, R. pedestris, N. lugens and C. kiangsu, O. infernalis, L. migratoria, C. nigricornis etc*. respectively (Figure: 2). The tree was constructed using maximum likelihood method so the results shows the related protein has a probability of distribution by maximizing a likelihood function.

**Figure 2:**
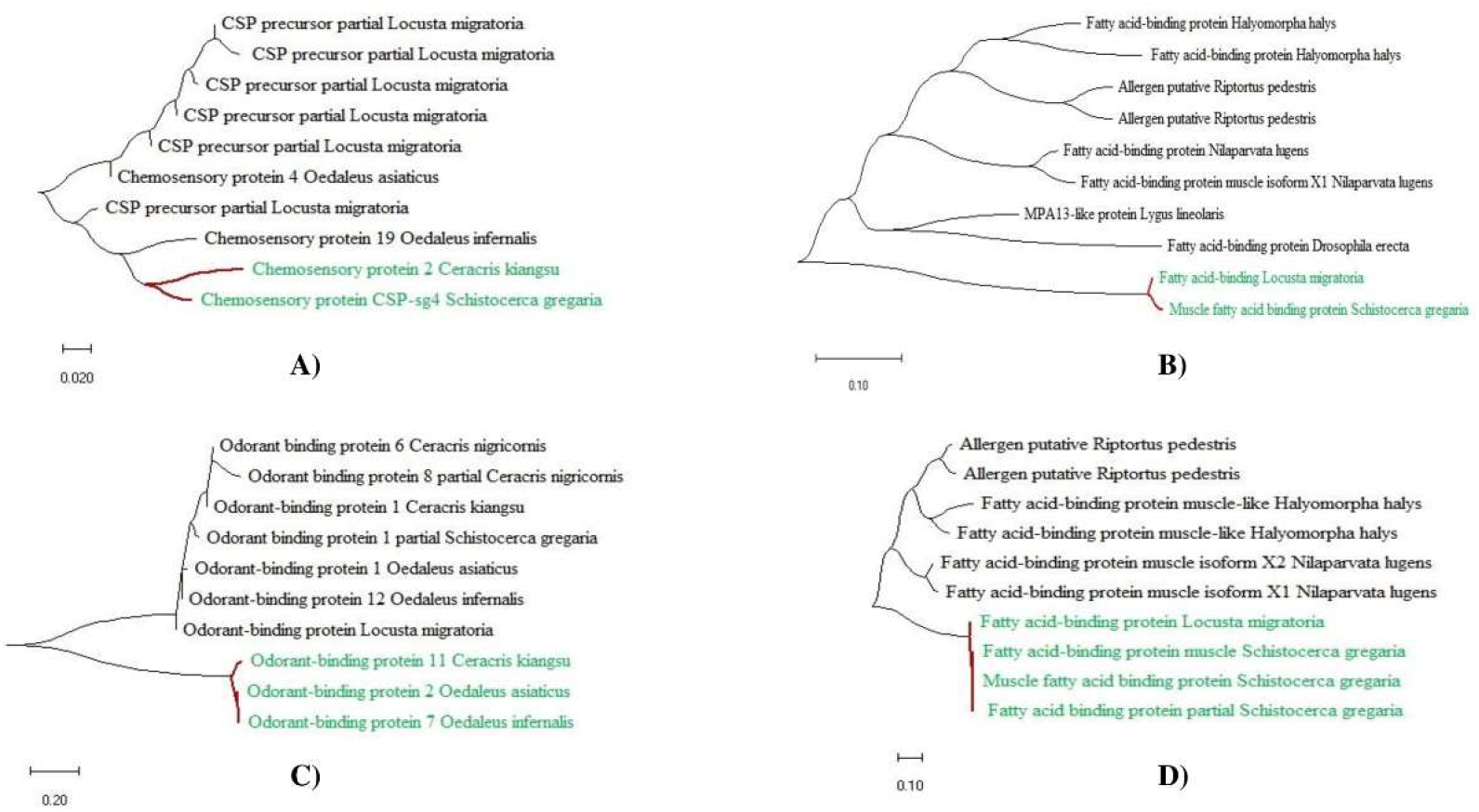
Phylogenic tree analysis; A) Lipid binding protein of *Schistocerca gregaria*; B) Muscle fatty acid binding protein of *Schistocerca gregaria*, C) Odorant binding protein of *ocusta igratoria*; D) Fatty acid binding protein of *Locusta migratoria*.

### 3.3. Molecular docking of locust protein/protein domain against pesticides

All of the retrieved structures of two different locust sp. (*S. gregaria and L. migratoria*) proteins/protein-domains (macromolecules) and pesticide (ligands) were prepared and employed for molecular docking to predict the affinity between above mentioned ligands and the macromolecules. The pesticides were ranked based on global binding energy and the results insights that top three scorers (pesticide) were similar for each of the macromolecules in terms of minimum binding energy (Supplementary File 2). In each case biphenthrin, diafenthiuron and silafluofen, showed best binding interactions with the four studied macromolecules (Figure 3 and Table 2). Moreover, silafluofen showed highest binding affinity with *L. migratoria* Fatty acid binding protein (−50.41 kcal/mol) (figure 4:B and Table 2) and also with Fatty-acid binding protein from *S. gregaria* (−48.73 kcal/mol) (figure 4:D and Table 2), while diaphenthiuron has bound with *L. migratoria* odorant binding proteins with a binding energy of −53.85 kcal/mol (Figure 4:C and Table 2) and biphenthrin bound with lipid binding protein from *S. gregaria* with a binding energy of −38.20 kcal/mol. (Figure 4A and Table 2). bendiocarb were used as a control in our study because of it’s uses for controlling locust problems worldwide [47].

**Figure 3.**
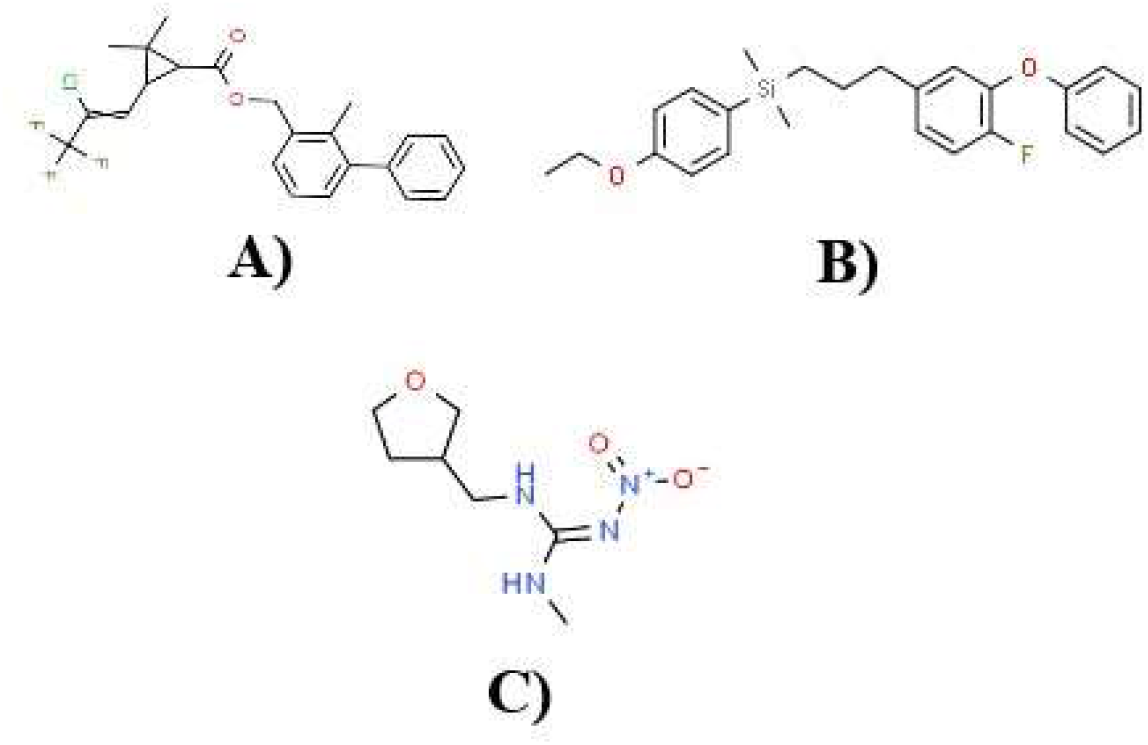
Chemical structure of (A)Biphenthrin, (B) Silafluofen and (C) Diafenthiuron.

**Figure 4:**
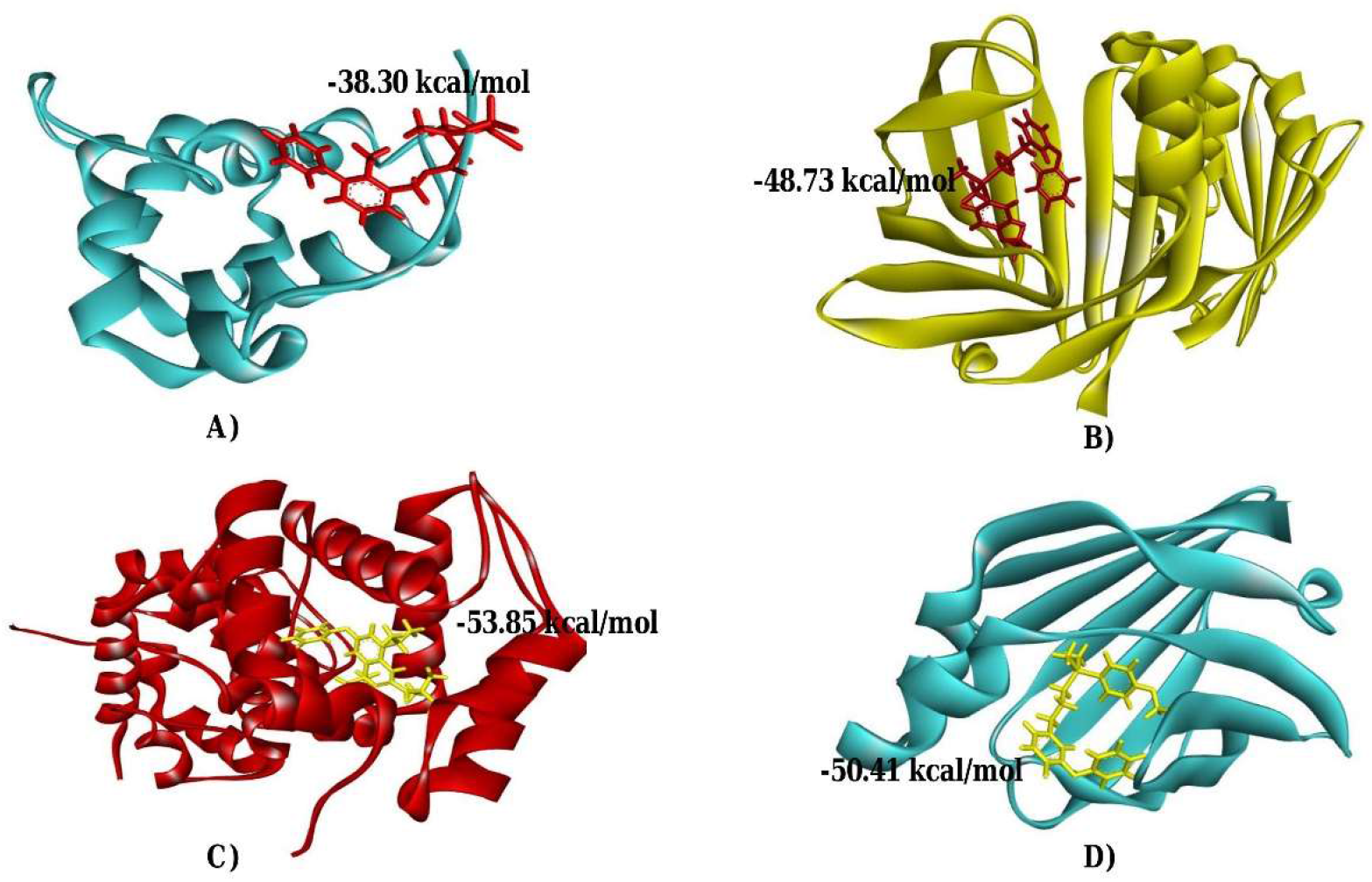
Molecular interaction of A) Biphenthrin with Lipid binding protein of *Schistocerca gregaria*; B) Silafluofen with Muscle fatty acid binding protein of *Schistocerca gregaria*, C) Diafenthiuron with Odorant binding protein of *Locusta migratoria*; D) Silafluofen with Fatty acid binding protein of *Locusta migratoria*.

**Table 2:** Analysis of Global binding energy and interaction sites of the screened top three pesticides.

| **Protein Name** | **Pesticides Name** | **Global Energy** | **Score** | **Area** | **ACE** | **Ligand binding residues** |
| --- | --- | --- | --- | --- | --- | --- |
| **2GVS** | Control (Bendiocarb) | -25.4 | 3110 | 368.2 | -4.28 | Leu13, Asp14, Lys70 |
|  | Biphenthrin | -38.20 | 5278 | 681.3 | -6.87 | Glu2, Lys3, Tyr4, Thr5, Thr6, Tyr8, Asp41 |
|  | Diafenthiuron | -33.99 | 4502 | 647.9 | -8.61 | Glu2, Lys3, Tyr4, Tyr8, Asp42, Glu44, Asn61, Lys63 |
|  | Silafluofen | -30.06 | 4798 | 617.6 | -7.43 | Lys3, Tyr4, Thr6, Tyr8, Ala40, Asp41, Glu44 |
| **1FTP** | Control (Bendiocarb) | -29.32 | 3710 | 428.7 | -7.44 | Lys10, Val40, Lys56, Ala58, Glu109, Val116, Gln133 |
|  | Biphenthrin | -42.46 | 5718 | 686.7 | -7.59 | Pro39, Ile41, Thr57, Lys60, Glu74, Leu77, His96, Gln98, Ile106, Ile117, Arg128, Tyr130 |
|  | Diafenthiuron | -42.02 | 5544 | 663.4 | -13.64 | Phe17, Ala33, Gly34, Pro39, Ser55, Lys60, Leu77 |
|  | Silafluofen | -48.73 | 5920 | 746.5 | -11.65 | Tyr20, Ile41, Thr54, Ser55, Thr62, Leu77, Gln98, Ile106, Ile119 |
| **4PT1** | Control (Bendiocarb) | -33 | 3464 | 437.8 | -10.49 | Tyr36, Phe47, Leu51, Ile54, His107, Tyr118, Val121, Val122, Phe125 |
|  | Biphenthrin | -47.05 | 5698 | 5698 | -13.24 | Ile11, Leu51, Ile54, Pro77, His107, Val121, Val122, |
|  | Diafenthiuron | -53.85 | 4690 | 696.1 | -19.34 | Ile11, Tyr50, Leu51, Ile54, Met55, Asn75, Val76, Pro77, His107, Tyr110, Tyr118, Val121, Val122, Phe125 |
|  | Silafluofen | -35.22 | 5106 | 686.2 | -8.52 | Arg35, Lys111, Pro115, Ser119 |
| **2FLJ** | Control (Bendiocarb) | -26.08 | 3720 | 461.3 | -6.93 | Phe17, Tyr20, Met21, Leu77, Ile106, Ile117, Ile119, Arg128 |
|  | Biphenthrin | -44.00 | 5862 | 770.3 | -7.36 | Tyr20, Leu37, Pro39, Lys60, Leu77, Gln98, Ile106, Ile117, Arg128 |
|  | Diafenthiuron | -45.62 | 5974 | 759.2 | -11.68 | Met21, Val26, Leu53, Phe64, Leu77, Ile106, Arg108, Arg128, Tyr130 |
|  | Silafluofen | -50.41 | 6100 | 846.6 | -12.65 | Phe17, Tyr20, Lys60, Thr62, Phe64, Leu77, Ile106, Arg108, Ile119 |

### 3.4. Analysis of pesticide inhibition surface hotspots

The molecular interaction between the ligands and target protein reveals the inhibition hotspots. It reveals the ligands were well fitted to the binding pockets of 2GVS by interacting amino acids like Tyr4, Tyr8, Ala 40 (Figure 5:A); 1FTP protein the ligand interacting amino acid residues such as Ile41, Leu77, Ile106, Ile117 etc. (Figure 5:B). The ligands attached with 4PT1 by interacting Ile54, Met55, Val76, Pro77, Tyr110, Tyr118, Val121, Val122 etc. (Figure 5:C). For 2FLJ the interacting molecule are Pro115, Tyr 20, Leu 77, Ile106 etc. (Figure 5:D). These binding sites can play a pivotal role for inhibiting the targeted proteins.

**Figure 5:**
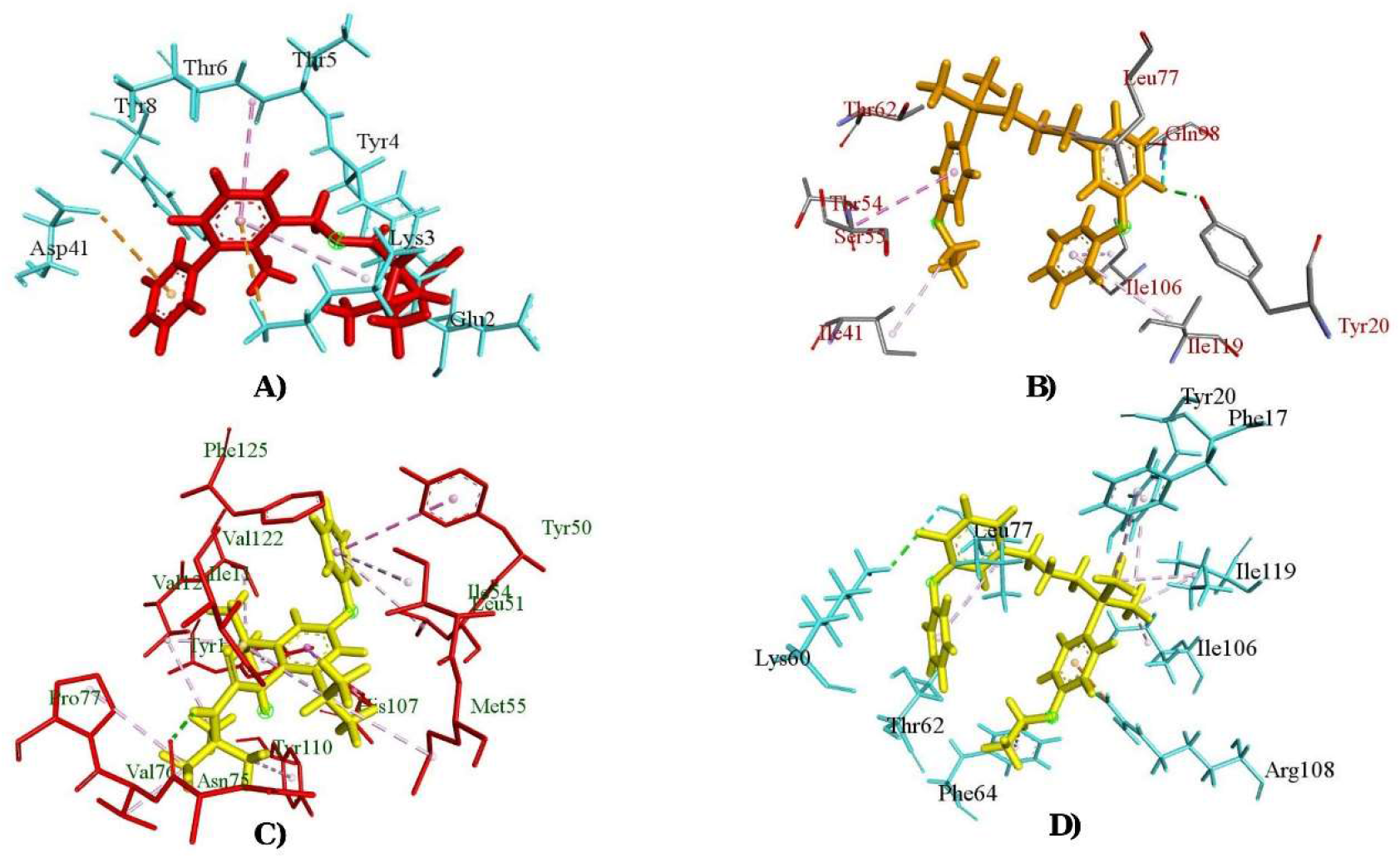
Ligand binding residues of A) Biphenthrin with Lipid binding protein of *Schistocerca gregaria*; B) Silafluofen with Muscle fatty acid binding protein of *Schistocerca gregaria*, C) Diafenthiuron with Odorant binding protein of *Locusta migratoria*; D) Silafluofen with Fatty acid binding protein of *Locusta migratoria*.

### 3.5. Free binding energy calculation and Molecular Dynamics stimulation

The free binding energy was calculated and found the pesticides had relatively good interactions with all the targeted proteins as evident from their negative binding free energies with biphenthrin and diafenthiuron having the highest binding affinity (Table 3), Notably, the 4PT1 protein with both biphenthrin and diafenthiuron showed maximum free binding energy among the studied complexes, with negative value describing the potential ease and stable interaction for the complex. For 4PT1-biphenthrin complex the MM/PBSA was calculated −27.38 kcal/mol and in MM/GBSA method it was −34.76 kcal/mol; 4PT1-diafenthiuron complex showed −24.77 kcal/mol and −34.08 kcal/mol for MM/PBSA and MM/GBSA, respectively.

**Table 3:** Binding free energy (kcal/mol) of the interaction of the selected pesticides with targeted protein.

| Macromolecules | Binding free energy (kcal/mol) | Biphenhrin | Diafenthuron |
| --- | --- | --- | --- |
| **2GVS** | MM/PBSA | -11.48 | -22.36 |
|  | MM/GBSA | -14.45 | -26.11 |
| **1FTP** | MM/PBSA | -7.16 | -6.70 |
|  | MM/GBSA | -16.85 | -14.82 |
| **4PT1** | MM/PBSA | -27.38 | -24.77 |
|  | MM/GBSA | -34.76 | -34.08 |
| **2FLJ** | MM/PBSA | -13.00 | -9.85 |
|  | MM/GBSA | -27.15 | -21.39 |

The most widely used molecular dynamics is the conventional type which describe the binding mode of ligand and the variation of protein internal dynamics. The normal mode analysis can be done by evaluating the deformability status of a complex, the experimental b factor and the eigen value.

The main-chain deformability which is a theoretical potential of any protein to deform at each of its residues. The results of our studied complexes revealed that all complexes had low flexibility potential and show resistance to deform (Figure 6: A-i, B-I, C-i, D-i; 7: A-i, B-i, C-i, D-i). The experimental B-factor which compares the normal mode analysis and the PDB field of the protein and predicted the corresponding uncertainty of their atomic positions showed that odorant binding protein of *L. migratoria* had the highest rigidity among the selected proteins and significance fluctuations weren’t found which proved the fact of lower loops (Figure 7:C-ii, D-ii). The energy required to deform the proteins is for all complexes and among studied complexes, the fatty acid binding protein from *L. migratoria* flight muscle and odorant binding protein of *L. migratoria*, as indicated by the highest eigenvalue of 6.227777 × 10^−4^, 3.065329 × 10^−5^ compared to muscle fatty-acid binding protein of *S. gregaria* and lipid binding protein of *S. gregaria* which eigen value of 9.914118 × 10^−5^, 8.045085 × 10^−5^ respectively (Figure 6:A-vi, B-vi, C-vi, D-vi; 7: C-vi, D-vi).

**Figure 6:**
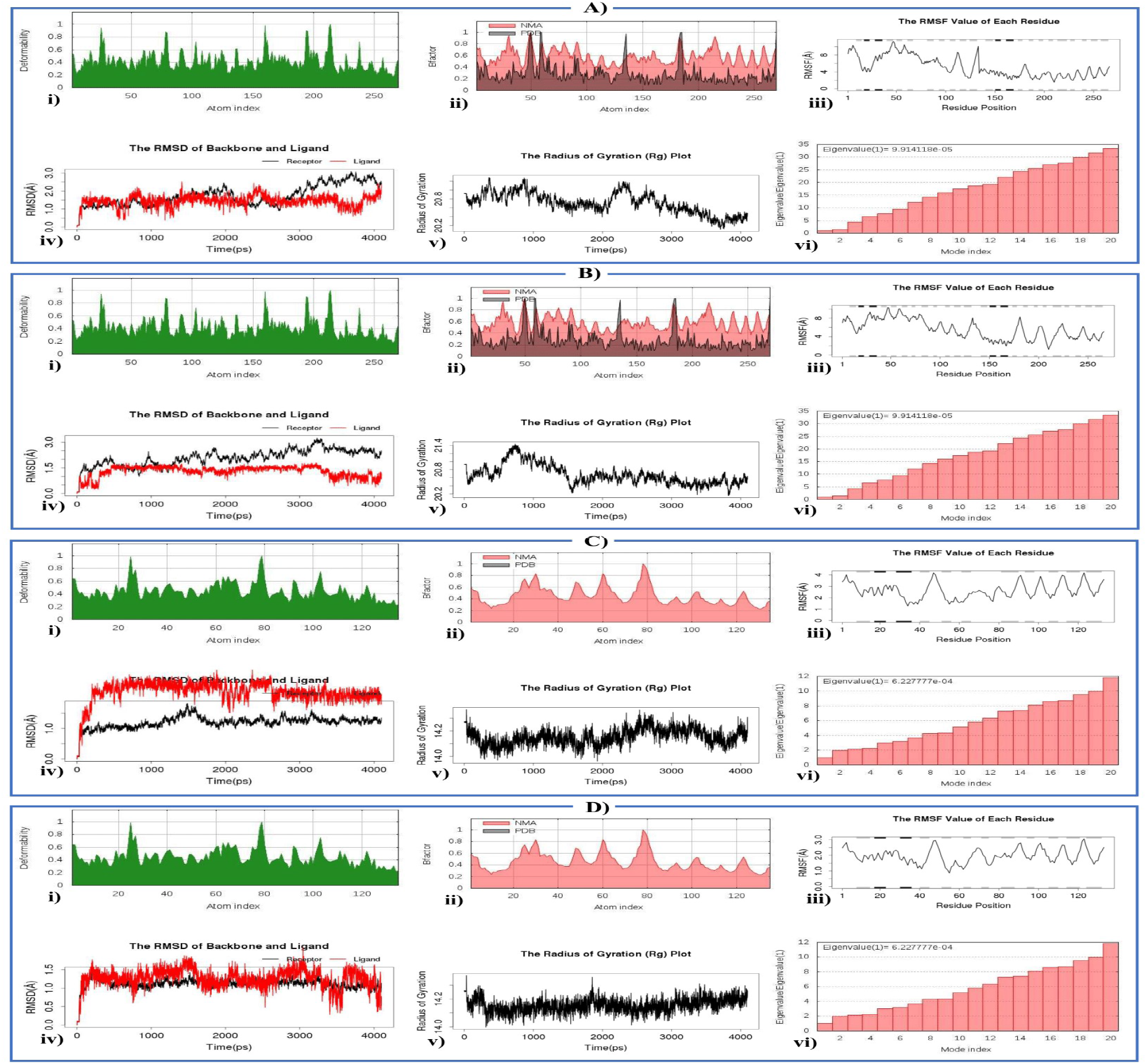
**Deformability analysis:** A-i) 1FTP with Biphenthrin, B-i) 1FTP with Diafenthiuron, C-i) 2FLJ with Biphenthrin, D-i) 2FLJ with Diafenthiuron; **B factor Analysis**: A-ii) 1FTP with Biphenthrin, B-ii) 1FTP with Diafenthiuron, C-ii) 2FLJ with Biphenthrin, D-ii) 2FLJ with Diafenthiuron; **RMSF plot**: A-iii) 1FTP with Biphenthrin, B-iii) 1FTP with Diafenthiuron, C-iii) 2FLJ with Biphenthrin, D-iii) 2FLJ with Diafenthiuron; **RMSD plot**: A-iv) 1FTP with Biphenthrin, B-iv) 1FTP with Diafenthiuron, C-iv) 2FLJ with Biphenthrin, D-iv) 2FLJ with Diafenthiuron; **Rg plot**: A-v) 1FTP with Biphenthrin, B-v) 1FTP with Diafenthiuron, C-v) 2FLJ with Biphenthrin, D-v) 2FLJ with Diafenthiuron; **Eigen value**: A-vi) 1FTP with Biphenthrin, B-vi) 1FTP with Diafenthiuron; C-vi) 2FLJ with Biphenthrin, D-vi) 2FLJ with Diafenthiuron.

**Figure 7:**
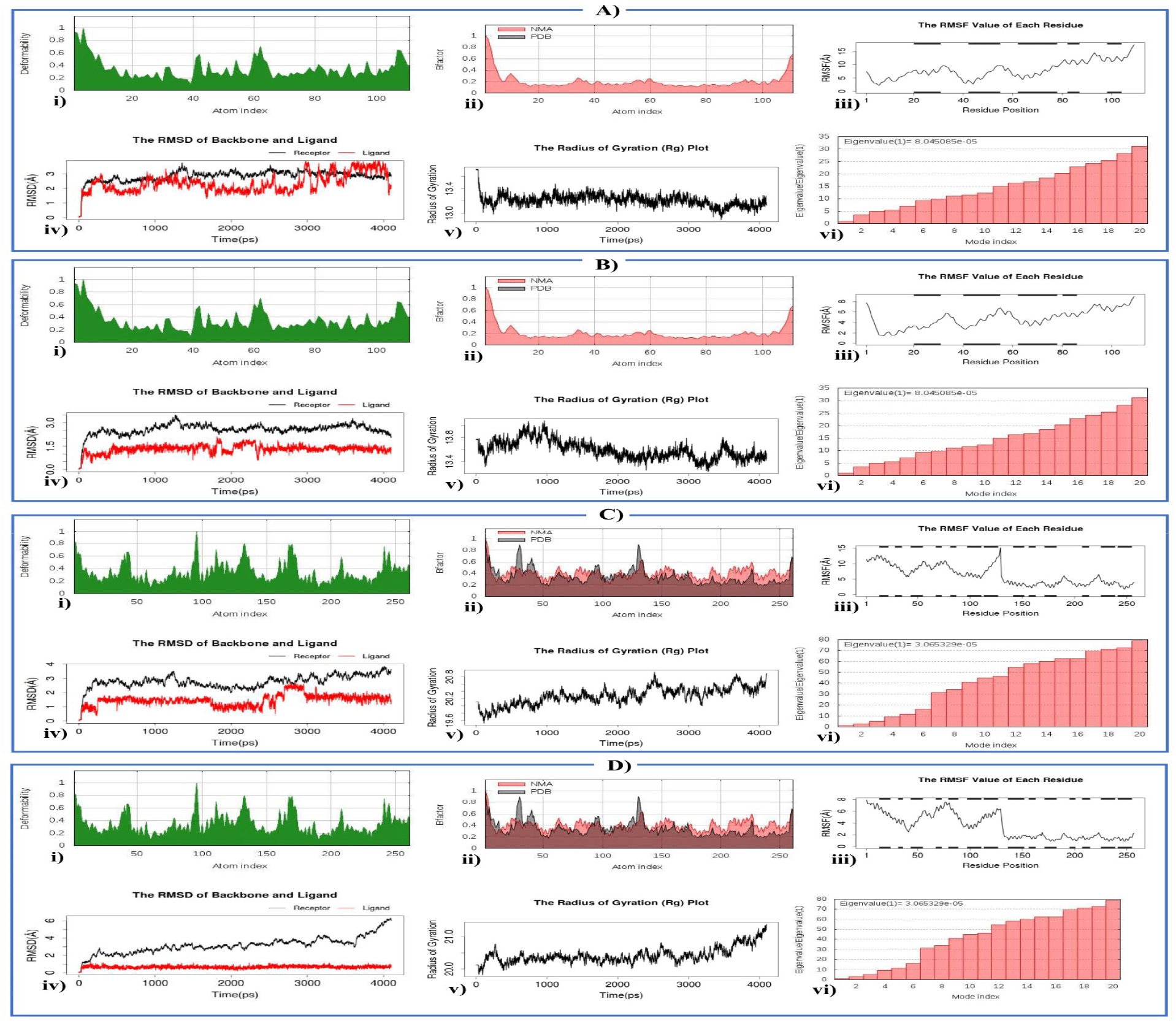
**Deformability analysis:** A-i) 2GVS with Biphenthrin, B-i) 2GVS with Diafenthiuron, C-i) 4PT1 with Biphenthrin, D-i) 4PT1 with Diafenthiuron; **B factor Analysis**: A-ii) 2GVS with Biphenthrin, B-ii) 2GVS with Diafenthiuron, C-ii) 4PT1 with Biphenthrin, D-ii) 4PT1 with Diafenthiuron; **RMSF plot**: A-iii) 2GVS with Biphenthrin, B-iii) 2GVS with Diafenthiuron, C-iii) 4PT1 with Biphenthrin, D-iii) 4PT1 with Diafenthiuron; **RMSD plot**: A-iv) 2GVS with Biphenthrin, B-iv) 2GVS with Diafenthiuron, C-iv) 4PT1 with Biphenthrin, D-iv) 4PT1 with Diafenthiuron; **Rg plot:** A-v) 2GVS with Biphenthrin, B-v) 2GVS with Diafenthiuron, C-v) 4PT1 with Biphenthrin, D-v) 4PT1 with Diafenthiuron; **Eigen value**: A-vi) 2GVS with Biphenthrin, B-vi) 2GVS with Diafenthiuron; C-vi) 4PT1 with Biphenthrin, D-vi) 4PT1 with Diafenthiuron.

From the result, the average RMSDs of CA atoms were normally distributed around 1-3 Å (Figure 6: A-iv, B-iv, C-iv, D-iv; 7:A-iv, B-iv, C-iv) except the odorant binding protein of *L. migratoria* - diafenthiuron complex which shows a bit fluctuation (Figure 7:D-iv). These results justified the true binding pose of the complexes. Although complexes of to muscle fatty-acid binding protein of *S. gregaria* - biphenthrin, fatty acid binding protein from *L. migratoria* flight muscle-diafenthiuron, lipid binding protein of *S. gregaria*-biphenthrin shows receptor-ligand equilibrium state (Figure 6: A-iv, D-iv; 7: A-iv). The Rg value range fluctuated between 13.4 to 21.4 for all complexes (Figure 6: A-v, B-v, C-v, D-v; 7: A-v, B-v, C-v, D-v). The RMSF plot of the complexes was found to align with all the target protein residues. The per residue Root Mean Square Fluctuations (RMSF) (Figure 6: A-iii, B-iii, C-iii, D-iii; 7: A-iii, B-iii, C-iii, D-iii) showed regular fluctuation pattern between 3 and 15 Å.

### 3.6. Toxicity analysis of pesticides

Prediction of various toxicity parameters such as AMES toxicity, minnow toxicity, fish toxicity, bird toxicity etc. were analyzed (Table 4). Estimated LD_50_ for biphenthrin, diafenthiuron and silafluofen were 2.776, 2.432 and 2.433 mol/kg, respectively. The fish toxicity results revealed the pLC50 for biphenthrin, diafenthiuron and silafluofen were −1.0555 0.3532 and 0.6658 mg/L respectively, means the silafluofen were tends to more toxic to fish than the other two pesticides. None of the compounds showed any undesired effects such as hERG I inhibitor, skin sensitisation, but silafluofen may possess some toxic consequences to bird and also exhibits positive result in AMES toxicity test.

**Table 4:** Toxicity pattern analysis of screened top three pesticides.

| Toxicity patterns of top three screened pesticides |  |  |  |  |
| --- | --- | --- | --- | --- |
| Parameters | Biphenhrin | Diafenthuron | Silafluofen | Remarks |
| AMES toxicity | No | No | Yes | Predicted |
| Max. tolerated dose (human) (log mg/kg/day) | 0.32 | 0.465 | 0.602 |  |
| hERG I inhibitor | No | No | No |  |
| Oral Rat Acute Toxicity (LD<sub>50</sub>) (mol/kg) | 2.776 | 2.432 | 2.433 |  |
| Oral Rat Chronic Toxicity (LOAEL) (log mg/kg) | 0.966 | 1.516 | 1.997 |  |
| Hepatotoxicity | No | Yes | Yes |  |
| Skin Sensitisation | No | No | No |  |
| *T. pyriformis* toxicity (log ug/L) | 0.474 | 0.439 | 0.347 |  |
| Minnow toxicity (log mM) | -2.017 | -1.163 | -3.548 |  |
| Bird Toxicity | No<sup>1</sup> | No<sup>1</sup> | Yes<sup>1</sup> | [21] |
| Rat Acute Toxicity (LD<sub>50</sub>) (mol/kg) | 3.89<sup>2</sup> | 2.269<sup>2</sup> | No available Data | [44] |
| Fish Toxicity (LC<sub>50</sub>) (ug/L) | < 100<sup>3</sup> | No available Data | No available Data | [45] |

### 3.7. Prediction of structural Analogs of pesticides

Ligand-based virtual screening was performed to predict small compounds from living organisms from mVOC v2.0 database. These living organisms derived small compounds which may provide a next sophisticated step towards the next generation pest management strategies. Dioctyl benzene-1,2-dicarboxylate (Pubchem CID 8346) also named dioctyl phthalate and dibutyl benzene-1,2-dicarboxylate (Pubchem CID 3026) also known as dibutyl phthalate derived from two bacterial species were found analogous to Biphenthrin with prediction score 0.47 and 0.42 respectively. Moreover, results revealed the similarity of 4-ethyl-2-methoxy-6-methylphenol (Pubchem CID 183540) and 3-propylphenol (Pubchem CID 69302) derived from two fungal species with Silafluofen with high prediction score (Table 5). The findings suggest that these could be potential pest management candidates against the devastating notorious pests, thus this requires further experimental trials for validation.

**Table 5:** Prediction of Microbial volatile compound from mVOC.

| Pesticide Name | Microbial Volatile Compound structural analog | Structural Analog Pubchem CID | Score | Source of Analog | Source Kingdom |
| --- | --- | --- | --- | --- | --- |
| Biphenthrin | Dioctyl benzene-1,2-dicarboxylate | 8346 | 0.47 | *Pseudomonas putida* | Bacteria |
|  | Dibutyl benzene-1,2-dicarboxylate | 3026 | 0.42 | *Carnobacterium divergens*<br>*Pseudomonas fragi* | Bacteria |
| Dinotefuran | 2,2,4,4-tetramethyloxolane | 137904 | 0.29 | *Klebsiella pneumoniae* | Bacteria |
| Silafluofen | 4-ethyl-2-methoxy-6-methylphenol | 183540 | 0.37 | *Tuber aestivum* and<br>*Tuber melanosporum* | Fungi |
|  | 3-propylphenol | 69302 | 0.36 |  | Fungi |

## 4. Discussion

The Desert Locust continues its footsteps, becoming a major threat to food security in several country of the world. Strategies of their invasion hold a high expense for the impacted nations, also a threat for ecological balance [6,48]. GIS based cartography illustrated the worldwide distribution of desert locust where *S. gregaria* and *L. migratoria* were the major desert locust species [6,7]. The results also indicate the swarm migration extend to Indian subcontinent and middle east in recent outbreak. It might be accelerated by the favorable environmental condition *viz*. temperature pattern, sudden rainfall sometimes direction of wind for locust migration [49,50]. Recently, almost thirty country has infested by this devasting pest and also threaten to several neighbor country [17]. Several pesticides were used widely in various countries to kill or control locust like bendiocarb but it doesn’t seem much more effective for controlling locusts [47,51]. Though some pesticides candidates are in the trial stages but already many of them raise controversial issues for their negative consequences towards environment and humans [52,53]. The world is intended to find an effective and justified controlling measures for minimizing the devastation by any possible means [54]. Several pesticides were evaluated against the proteins of desert locust using computational approaches to find out the best possible mean with higher effectivity and less possible environmental and human impact.

The computer aided inhibitor design has been contributed to accelerate the inhibitor discovery for several pest problems. By these approaches, the possibilities of potent small molecules as ligands/inhibitors can be explored [55]. Pesticides like chlorpyrifos, deltamethrin, diflubenzuron, lambda-cyhalothrin has been used as potential locust controlling agent. But these pesticides don’t pass the expectation level. They have also a wide range of toxic complications [47,56]. The several studies showed Bendiocarb used against locust that has similar level of activity as previously mentioned locust controlling pesticides with comparatively higher level of toxicity [47,51]. We used bendiocarb as a positive control in our study for our targeted proteins.

Some commonly pesticides were screened against desert locust species proteins such as lipid binding protein (2GVS) and muscle fatty acid binding protein (1FTP) from *S. gregaria* along with odorant binding protein (4PT1) and flight muscle fatty acid binding protein (2FLJ) from *L. migratoria* using a molecular docking approach [57].

The lipid binding protein (2GVS) of *S. gregaria* is a chemosensory protein. Chemosensory proteins are soluble proteins found in insect’s chemosensory organs which are responsible for chemical message transmission and this chemical stimulus has been referred extremely important for the survival of various insect species [58]. The muscle fatty acid binding protein (1FTP) of *S. gregaria* has a vital role in fatty acid transport and metabolism [59]. The odorant binding protein (4PT1) of *L. migratoria* is crucial to find their plant hosts through which they find feed for survival utilizing their olfactory system [60] and the flight muscle fatty acid binding protein (2FLJ) of *L. migratoria* plays a vital role in their wing movement by transportation of fatty acid [61].

Remarkably, three pesticides i.e. biphenthrin, diaphenthiuron and silaflufen scored best for each four macromolecules and bound with minimum global binding energy (Table 2) and Supplementary File 2). Silafluofen showed highest binding affinity with both *L. migratoria* fatty acid binding protein (−50.41 kcal/mol) (Figure 4B Table 2) and also with fatty-acid binding protein from *S. gregaria* (−48.73 kcal/mol) (Figure 4D Table 2), while diaphenthiuron has bound with *L. migratoria* odorant binding proteins with a binding energy of −53.85 kcal/mol (Figure 4C Table 2) and biphenthrin bound with lipid binding protein from *S. gregaria* with a binding energy of −38.20 kcal/mol. (Figure 4A Table 2). The scores of top candidates were higher than bendiocarb, a positive control used in the present study (Table 2). The silaflufen and biphenthrin generally acts in sodium channel modulators of pest and make a significant impact on their nerve, the other one which is diafenthiuron targets mitochondrial ATP synthase and seriously effects energy metabolism (Table 1) [62].

The molecular interaction between the ligands and target protein reveals the inhibition hotspots. It reveals the ligands were well fitted to the binding pockets of targeted proteins by interacting several amino acids like Tyr4, Tyr8, Ile119,Ile11, Tyr50, Leu51, Ile54, Met55 etc. by several kinds of bonding and most of bindings are relatively in hydrophobic environment, which may help to stabilize its conformation and these ligand-protein hydrophobic interactions can play a crucial role because these may help in stabilizing the complex and improved the efficacy of the inhibitor [63].

Molecular dynamics stimulation of biphenthrin and diafenthiuron was studied with all four protein complexes. The free binding energy calculation shows negative result for all proteins means the strong binding affinity and the normal mode analysis uncover that the complexes were stable with the ligand. The all docked complexes B factor and deformability analysis showed less hinges means the rigidity and stability of the complexes. The comparatively higher eigen value for locust flight muscle protein with pesticides complexes confirms its strong stiffness than others. The odorant binding protein of *L. migratoria* - diafenthiuron complex which shows a bit fluctuation (Figure 7:D-iv). The lower the RMSD distribution in Å, higher possibility of complex true pose. The founded results justified the true binding pose of the complexes. Notable, complexes of to muscle fatty-acid binding protein of *S. gregaria* - biphenthrin, fatty acid binding protein from *L. migratoria* flight muscle-diafenthiuron, lipid binding protein of *S. gregaria*-biphenthrin shows receptor-ligand equilibrium state ((Figure 6: A-iv,D-iv; 7: A-iv)). The Rg value range fluctuated between 13.4 to 21.4 for all complexes ((Figure 6: A-v, B-v, C-v, D-v; 7: A-v, B-v, C-v, D-v). The per residue Root Mean Square Fluctuations (RMSF) (Figure 6: A-iii, B-iii, C-iii, D-iii; 7: A-iii, B-iii,C-iii, D-iii)) showed regular fluctuation pattern. These analysis insights negligible chance of deformability as hinges are in the chain was not significant and strongly validate our prediction. According to our encouraging results, the toxicity pattern among these top three candidates reveals that biphenthrin and diafenthiuron were relatively safer than silaflufen in terms of bird, fish, human and other toxicity parameter (Table 4). The phylogenetic analysis of our targeted 4 protein from 2 different species shows close relationship with other pest species precisely to the various locusts such as *H. haly, R. pedestris, N. lugens, C. kiangsu, O. infernalis, R. pedestris, C. nigricornis.* We suggest, our top screened three pesticides may also prove their effectiveness against these pest species also. The structural similarity search of our top screened three pesticides found analogs to the several microbial volatile compounds. Dioctyl benzene-1,2-dicarboxylate (Pubchem CID 8346) and dibutyl benzene-1,2-dicarboxylate (Pubchem CID 3026) derived from two bacterial species were found analogous to biphenthrin with high prediction score 0.47 and 0.42 respectively (Table 5). Moreover, results revealed the similarity of 4-ethyl-2-methoxy-6-methylphenol (Pubchem CID 183540) and 3-propylphenol (Pubchem CID 69302) derived from two fungal species with silafluofen with high prediction score (Table 5). The findings suggested that these could be potential pest management candidates also against the devastating notorious locust, as they are derived from living microorganisms. They have the potential to break down the ice of the controversial argument with use of pesticides which requires further experimental field trials for validation.

## 5. Conclusion

The result suggests that biphenthrin, diafenthiuron and silaflufen could be option to combat desert locust. Furthermore, two biological derived structural analogs from mVOC i.e. dioctyl phthalate and dibutyl phthalate may be effective and show inhibitory properties against the desert locust essential proteins. Due to the encouraging results, we highly recommend further in the field trials for the experimental validation of our findings.

## Acknowledgment

We acknowledge Department of Entomology, Sylhet Agricultural University and Department of Plant and Environmental Biotechnology, Sylhet Agricultural University for their technical support.

